# Effect of antiretroviral protease inhibitors on *Plasmodium falciparum* erythrocyte egress and invasion

**DOI:** 10.1101/2023.10.05.560994

**Authors:** Danny W. Wilson, Sonja Frolich, Katherine T. Andrews, Tina S. Skinner-Adams

## Abstract

**Background:** Anti-retroviral protease inhibitors directly inhibit the growth of asexual blood stage malaria parasites, however, this activity is not fully understood. While mode of action hypotheses have included parasite aspartic protease (plasmepsin) inhibition, current data suggest that digestive vacuole plasmepsins I-IV are not essential for asexual parasite survival, that plasmepsins VI-VIII are not expressed in these parasites and that antiretroviral protease inhibitors are poor inhibitors of plasmepsin V. The remaining plasmepsins, IX and X, have recently been shown to be essential for merozoite egress and invasion, playing important roles in the processing of key proteins including the rhoptry bulb protein *Pf*RAP1, and subtilisin-like serine protease *Pf*SUB1, respectively. To further understand the antiplasmodial activity of antiretroviral protease inhibitors, here we investigated the impact of tipranavir, lopinavir, ritonavir and saquinavir on the processing of *Pf*RAP1, the *Pf*SUB1-processed *Pf*MSP1, and the egress and invasion of *P. falciparum* parasites from human erythrocytes.

**Methods:** The effect of tipranavir, lopinavir, ritonavir and saquinavir on *P. falciparum* parasite egress and invasion was assessed using synchronized asexual blood stage *P. falciparum* parasites. Schizont rupture and purified merozoite invasion were performed with and without drug and quantified by flow cytometry analysis. The impact of selected antitretroviral protease inhibitors on *Pf*RAP1 and *Pf*MSP1 processing was assessed by Wesstern blot.

**Results:** The effect of tipranavir, lopinavir, ritonavir and saquinavir on the egress and invasion of *P. falciparum* parasites from human erythrocytes varied considerably, but was low at concentrations shown to inhibit *P. falciparum* asexual parasite growth *in vitro* and negligible at clinically relevant concentrations. While the treatment of parasites with the antiretrovial protease inhibitors appeared to reduce the overall expression of *Pf*RAP1 and *Pf*MSP1, the processing of these proteins was not inhibited by concentrations known to inhibit parasite growth *in vitro*.

**Conclusions:** The limited activity of tipranavir, lopinavir, ritonavir and saquinavir on the egress and invasion of *P. falciparum* parasites from human erythrocytes and the processing of *Pf*RAP1 and *Pf*MSP1 suggests that plasmepsin IX and X are unlikely to be the primary targets of these drugs in these parasites.

## Background

Antiretroviral protease inhibitors kill malaria parasites *in vitro* [1, 2], *ex vivo* and *in vivo* [3, 4] at concentrations that are clinically achieved in patients receiving these drugs for the treatment of HIV infection. While the relatively high concentrations of each drug required for antiplasmodial activity (μM) precludes their use as first-line antimalarial candidates, data showing that they are active against multiple *Plasmodium* species [3, 4] at various stages of parasite development [1, 4-6] and suggest that an understanding of the mode of action of these drugs may aid the development of specific and more effective chemotherapeutics.

While glucose transport has been postulated as the anti-*Plasmodium* target of the antiretroviral protease inhibitor lopinavir, the concentrations of lopinavir required for glucose transport inhibition in parasites (IC_50_ 16 μM) are not in-line with those required for growth inhibition (IC_50_ 1.9 μM), suggesting additional anti-parasitic targets are likely at play [7]. As antiretroviral protease inhibitors are potent inhibitors of HIV aspartyl protease, it has long been hypothesised that they target a malaria parasite aspartic protease [8, 9]. There are ten aspartyl proteases, collectively known as plasmepsins, in *P. falciparum* [10]. However three of these (plasmepsins IV-VIII) are not expressed in asexual stages of parasite development (plasmodb.org). Transgenic parasite data has also demonstrated that the four digestive vacuole plasmepsins (plasmepsins I, II, III (HAP) and IV) are not essential to asexual parasite growth *in vitro* and that loss of these plasmepsins does not impact sensitivity to antiretroviral protease inhibitors [9]. The remaining three enzymes, plasmepsins V, IX and X, are essential to asexual erythrocytic-stage parasite development, however, recombinant protein inhibition data suggest that antiretroviral protease inhibitors have limited activity against plasmepsin V [11]. Thus, plasmepsins IX and X could be considered likely targets of the antiretroviral aspartic protease inhibitors in *Plasmodium* parasites. Plasmepsin IX and X are essential maturases that are required for processes including merozoite egress and invasion [12, 13]. While plasmepsin IX is involved in the processing of proteins including rhoptry protein 1 (*Pf*RAP1) and is essential for merozoite invasion, plasmepsin X has been shown to process multiple proteins including apical membrane antigen 1 (*Pf*AMA1) and the serine protease *Pf*SUB1, and to be essential for *P. falciparum* erthyrocyte egress and invasion [13]. During egress, *Pf*SUB1 has been shown to process multiple additional proteins including the merozoite surface protein *Pf*MSP1 [13]. To further investigate the role of plasmepsins IX and X in mediating the antiplasmodial activity of the antiretrivial protease inhibitiors, in this study, we investigated whether the drugs tipranvir, lopinavir, ritonavir and saquinavir inhibit *P. falciparum* erythrocyte egress and invasion. The processing of *Pf*RAP1 and *Pf*MSP1 in the presence of selected antiretrovial protease inhibitors was also investigated.

## Methods

Tipranavir was obtained from the AIDS Reagent Program, Division of AIDS, NIAID, NIH. All remaining antiretroviral protease inhibitors were obtained from Sigma-Aldrich. The effect of lopinavir, saquinavir, ritonavir and tipranavir on *P. falciparum* parasite egress and invasion was assessed using D10-*Pf*PHG parasites [14]. Parasites were cultured in human O^+^ erythrocytes, as previously described [14], and synchronised to a 4 h window (for schizont exposure experiments) or 6 h window (for merozoite invasion experiments) using sterile injectable heparin (Pharmacia) [15]. For schizont rupture assays, 45 μl of synchronised late schizont stage parasites (∼46 h post invasion for short treatment assays or ∼42 h post invasion for long treatment assays) were added to a 96-well round bottom plate at 2% parasitaemia and 1% haematocrit with 5 μl of drug or control (media or vehicle). Parasites were incubated with drug for 8 or 12 h (short or long assays, respectively) and stained with 5 μg/ml ethidium bromide (EtBr, Bio-Rad) for 10 min followed by immediate flow-cytometry analysis (Becton Dickinson, LSR), as previously described [16, 17]. For merozoite invasion inhibition assays, purified free merozoites (22.5 μl) [15], were mixed with 2.5 μl of drug or control in a 96-well round bottom plate for 10 min at 37°C before the addition of uninfected red blood cells to a final haematocrit of 0.5%. Plates were agitated (400 rpm) for 10 min and invasion was assessed within 1 h by flow cytometry. Heparin (2,000 IU) was included as a positive control in all assays. For asexual parasite IC_50_ determinations synchronised *P. falciparum* ring stage parasites (∼0-12 h post invasion) were exposed to compounds or controls in 96-well round bottom plates (1% parasitaemia and 1% haematocrit) for 72 h prior to staining with 10 μg/ml ethidium bromide for 10 min, washing and resuspension in fresh PBS. Parasite counts were determined by flow-cytometry and IC_50_ values were calculated using regression curve fit GraphPad Prism. All experiments were repeated on three separate occasions.

For Western blots, synchronous schizont-stage cultures (∼42 h post invasion) at 5-7% parasitaemia were treated with protease inhibitors, (tipranavir 100 μM or 50 μM; lopinavir 5 μM or 2.5 μM; ritonavir 100 μM or 50 μM; saquinavir 40 μM or 20 μM) for 6 h and lysed in 0.15% w/v saponin (ThermoFisher Scientific) for 10 min on ice. Pellets were washed in 0.075% w/v saponin and PBS until the supernatant ran clear. Saponin pellets were resuspended in reducing sample buffer (0.125 M Tris-HCl pH 7, 20% v/v glycerol, 4% v/v SDS, 10% v/v β-mercaptoethanol (Sigma-Aldrich), 0.002% w/v bromophenol blue (Sigma-Aldrich) and electrophoresed on 4-12% SDS-PAGE Bis-Tris gels (Bolt, Invitrogen) at 110 V for 70 min. Protein was transferred onto nitrocellulose membranes (iBlot, Invitrogen) at 20 V for 7 min. Membranes were blocked in 1% w/v skim milk with 0.05% v/v Tween 20 (Sigma-Aldrich) in PBS (1% milk PBS-T) for 1 h before incubation with the primary antibodies (rabbit anti-*Pf*EXP2 1/10000, rabbit anti-*Pf*MSP1-19 1/10000, mouse anti-*Pf*RAP1 1/10000) overnight at 4°C. Membranes were washed three times in PBS-T and then incubated for 1 h with secondary antibodies (IRDye 800 goat anti-mouse and IRDye 680 goat anti-rabbit (LI-COR Biosciences)) in milk PBS-T. Membranes were washed three times in PBS-T and once in PBS before being dried and imaged using an Odyssey Infrared Imaging System (LI-COR Biosciences).

## Results

The ability of tipranavir, lopinavir, ritonavir and saquinavir to inhibit the invasion of erythrocytes by free *P. falciparum* merozoites was assessed directly in assays using purified merozoites [15]. The only antiretroviral protease inhibitor to demonstrate a clear dose-dependent effect on merozoite invasion in this work was lopinavir (asexual parasite IC_50_ 3 μM), with invasion inhibitory values of 18%, 38% and 73% at 10 μM, 20 μM and 50 μM, respectively (**Figure 1**). While tipranavir and saquinavir (asexual parasite IC_50_ values 68 ± 18 μM and 26 ± 3 μM respectively) demonstrated similar activity to lopinavir at 50 μM (75% and 85% inhibition, respectively) this activity was not dose dependent. Interestingly, while the invasion inhibitory activities of lopinavir, ritonavir (asexual parasite IC_50_ 33 ± 3 μM) and saquinavir were negligible at clinically relevant concentrations (Cmax 15.3, 11.2 and 3.1 μM, respectively [3]), tipranavir demonstrated 75% invasion inhibiton at a clinically relevant concentration (**Figure 1A**; Cmax 60 to 185 μM [18]).

**Figure 1:**
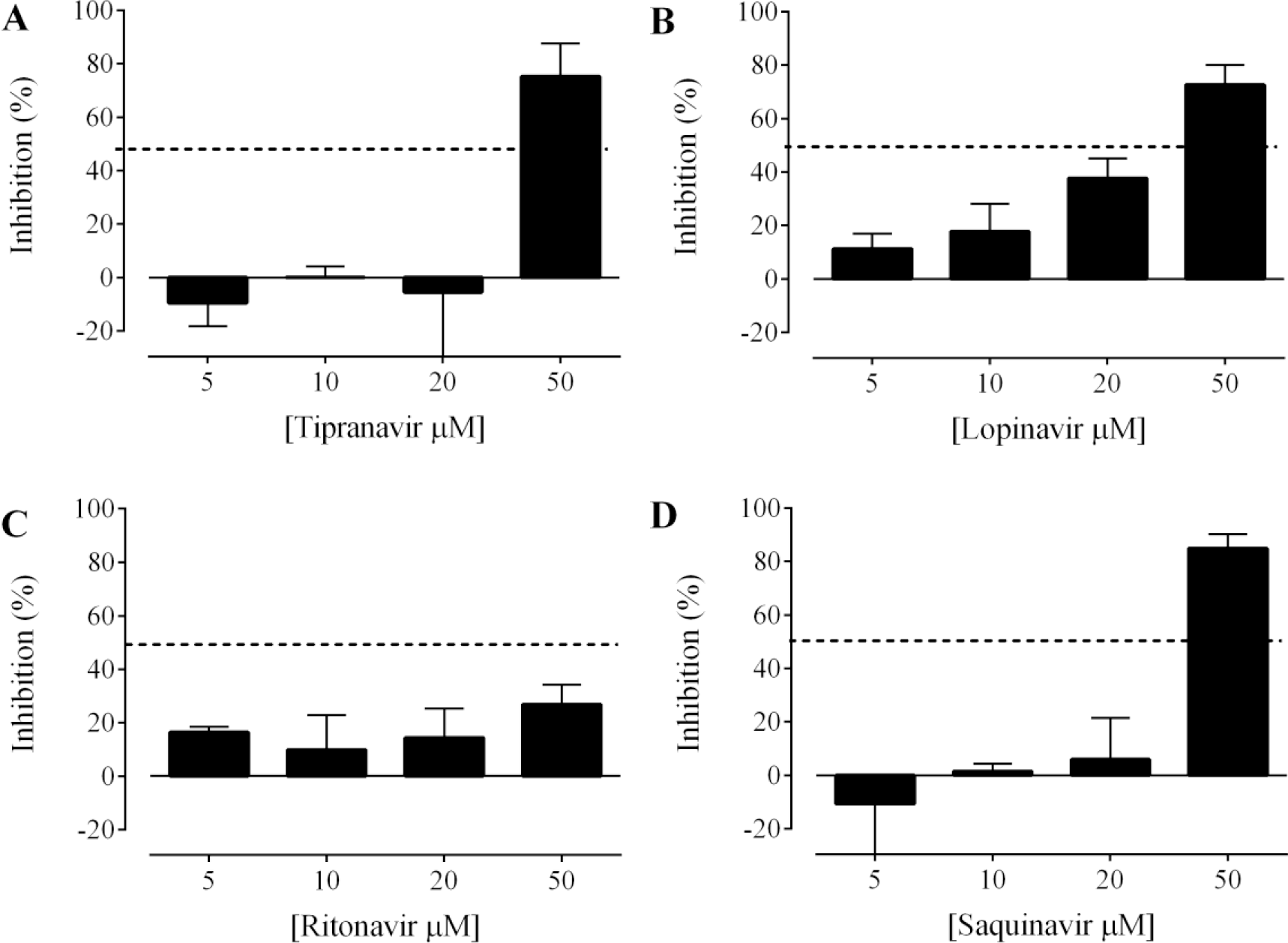
Inhibitory activity of antiretroviral protease inhibitors on the invasion of *P. falciparum* merozoites into human erythrocytes. *P. falciparum* D10-*Pf*PHG purified free merozoites were exposed to tipranavir (**A**), lopinavir (**B**), ritonavir (**C**) or saquinavir (**D**) for 10 min prior to the addition of fresh uninfected red blood cells to allow invasion. Following a further 10 min incubation, invasion was assessed by flow cytometry. Data are presented as mean (± SEM) % inhibition of ring-stage parasites for three independent experiments, each carried out in duplicate wells. Heparin (2,000 IU) was included as an invasion inhibitory control and inhibited invasion in all cases (mean % inhibition (± SEM) ring-stage parasites of 91.6 ± 3.7; not shown in figure). Dashed lines represent 50% inhibition.

As merozoite treatment assays are time restricted, due to the short window of merozoite invasion and viability, and the impact of plasmepsin IX and X inhibtion on protein processing and downstream egress and invasion has been shown to be time dependent [12, 13], we next sought to assess whether preincubation of developing merozoites within infected erythrocytes with selected antiretroviral protease inhibitors could impact egress and invasion. To do this we treated schizont stage parasites at ∼46 h post invasion for 8 h, or schizont stage parasites at ∼42 h post invasion for 12 h. Data derived from 8 h exposure experiments demonstrated that the treatment of schizonts with lopinavir resulted in a dose dependent reduction in ring stage parasites and a reduction of free merozoites. An ∼20% accumulation of schizonts was also seen at the highest treatment concentration (**Figure 2B**). These observations suggest the inhibition of schizont development or merozoite egress, rather than invasion inhibition. However the concentrations of lopinavir required for these effects was higher than clinically relevant levels (Cmax 15.3 μM [3]) and >20-fold above the concentration required to cause *in vitro* asexual stage growth inhibiton in longer-term (48-72 h) assays (IC_50_ 1-3 μM [3]; **Figure 1B**). Similarly, the reduced invasion associated with tipranavir (**Figure 2A**), ritonavir (**Figure 2C**), and saquinavir (**Figure 2D**) treatment was associated with a loss of free merozoites, an observation that is typically associated with the the inhibition of schizont rupture and merozoite egress. However, these phenotypes were typically observed above clinically relevant concentrations (**Figure 2**). These data suggest that treatment of parasites with antiretroviral protease inhibitors during the final 2 h of schizont development does not inhibit parasite invasion directly, but likely impacts schizont rupture at higher concentrations.

**Figure 2:**
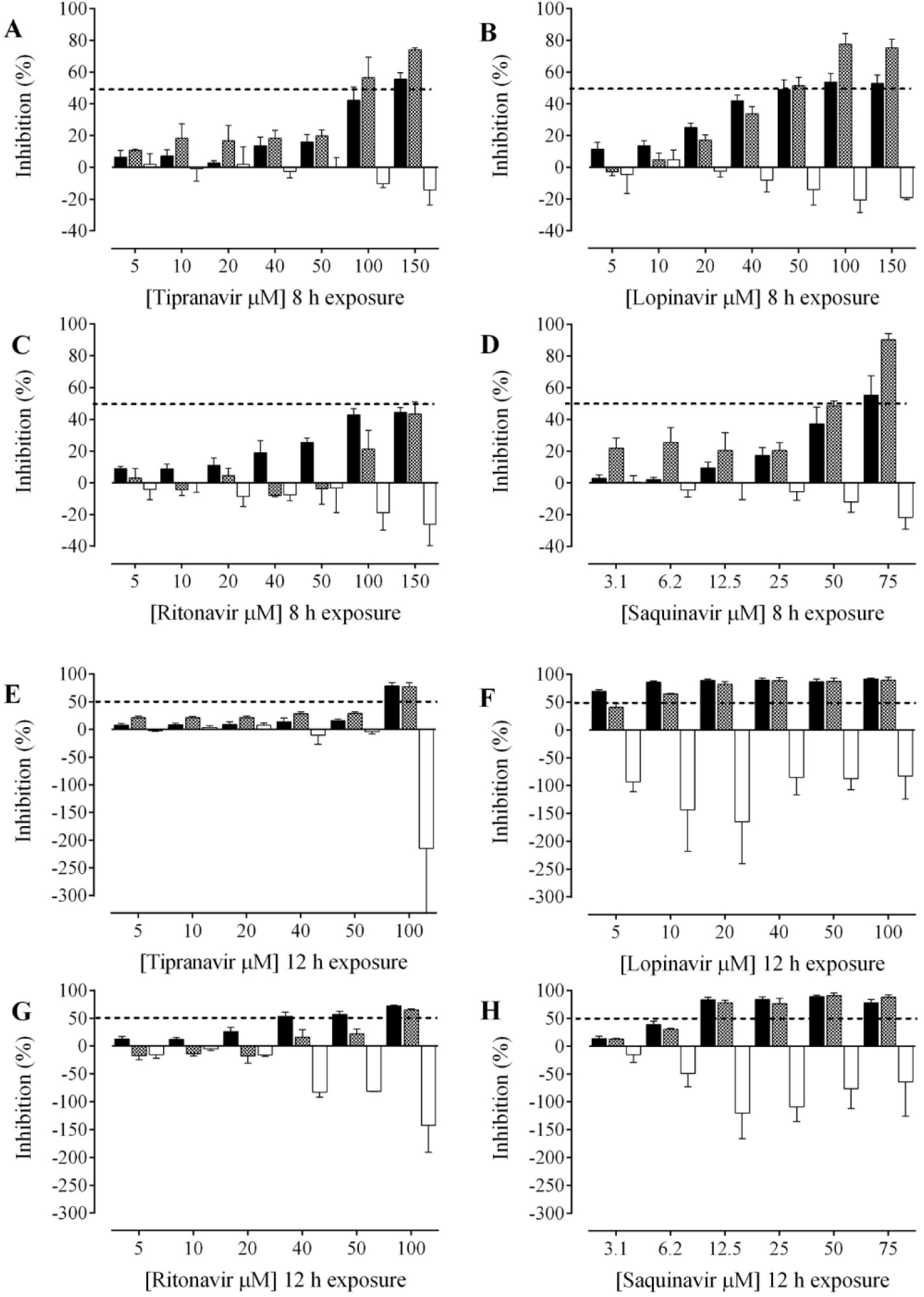
Activity of antiretroviral protease inhibitors on the egress and invasion of *P. falciparum* parasites from, and into, human erythrocytes. Synchronous schizont-stage *P. falciparum* D10-*Pf*PHG parasitized erythrocytes (∼46 h (**A-D**) or 42 h (**E-H**) post invasion) were exposed to tipranavir (**A & E**), lopinavir (**B & F**), ritonavir **(C & G**) or saquinavir (**D & H**) for 8 (**A-D**) or 12 h (**E-H**) prior to flow-cytometry analysis of schizont rupture and merozoite egress and invasion. Data are presented as mean (± SEM) % inhibition of ring (black), merozoite (hatched lines) and schizont (white) parasites of three independent experiments, each carried out in duplicate wells. Heparin (2,000 IU) was included as an invasion inhibitory control and inhibited invasion in all cases (mean inhibition (% ± SEM) of rings, merozoites and schizonts stage parasites of 85.6 ± 5.4, -22.7 ± 22.6 and -1.6 ± 17.9 respectively). Dashed lines represent 50% inhibition.

Since the essential proteolytic processing events mediated by plasmepsin IX and X may occur earlier in schizont development, we also tested the effect of a longer duration protease inhibitor treatment on synchronized schizonts (∼42 h post invasion; 48 h asexual lifecycle; ∼6 h prior to schizont rupture). Similar inhibitory profiles were observed for these long treatment experiments as for the shorter (∼46 h post invasion) assays. However, schizont rupture and merozoite egress inhibition was more pronounced for all treatments (**Figure 2E-H**). There was also a notable inverse correlation with the inhibition of ring stages and free merozoites and the presence of schizonts in these long duration treatments. However, this phenotype was most noticeable at treatment concentrations that were generally higher than the concentrations needed to kill asexual parasites (**Figure 2**). To assess whether the schizont accumlation seen in treatments was due to merozoite egress inhibition or direct schizont development inhibition, Giemsa stained thin smears of parasites treated with tipranvir, lopinavir, ritonavir and saquinavir at concentrations where loss of merozoite invasion was evident with extended schizont treatment (**Figure 2E-H**) were prepared. Unhealthy schizonts were observed in all treatments (**Figure 3**), indicating that the inhibition seen in egress and invasion assays was likely the result of schizont development inhibition rather than merozoite egress or invasion inhibition.

**Figure 3:**
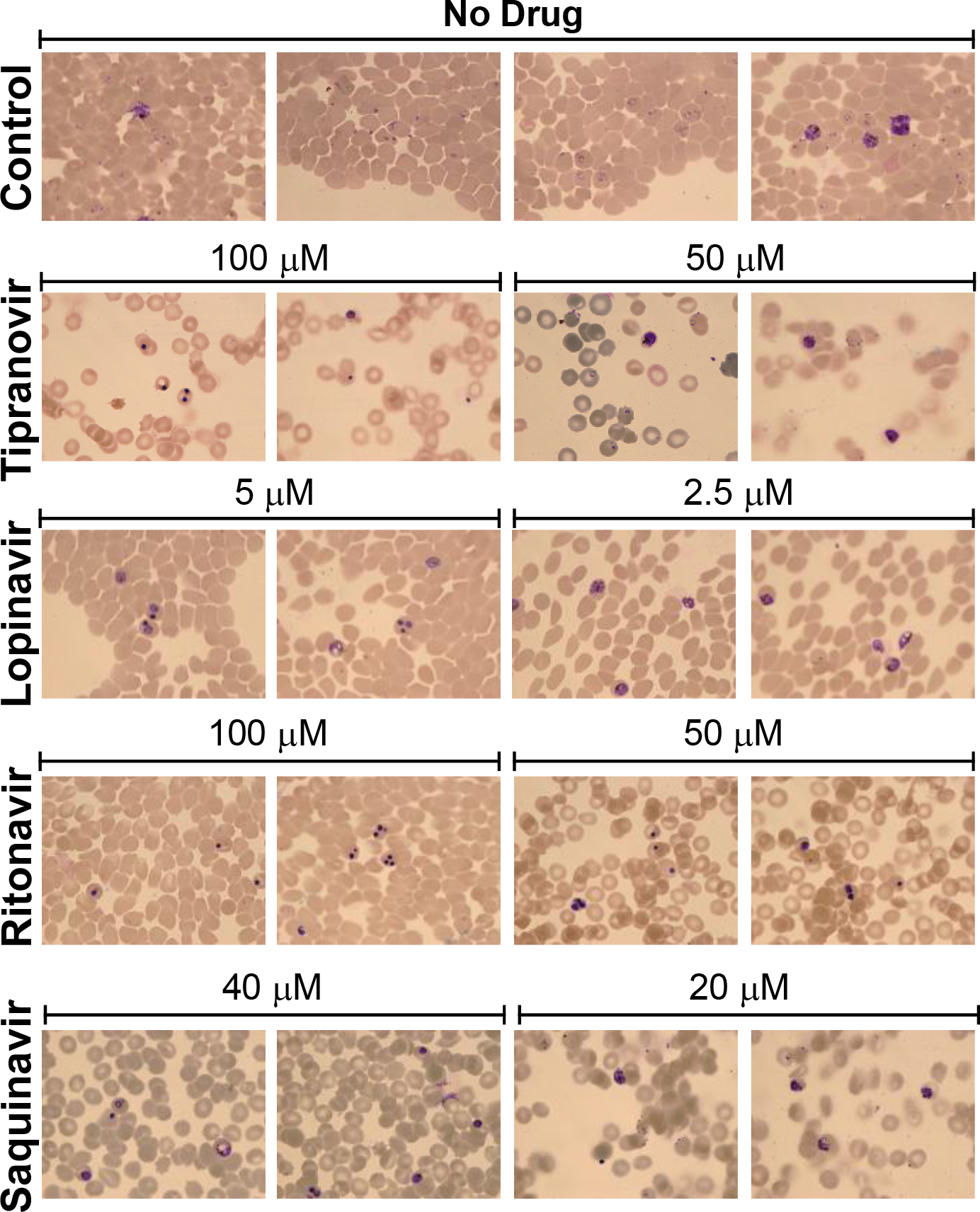
Imapct of antiretrovial protease inhibitors on parasite morphology. Synchronous schizont-stage *P. falciparum* D10-PfPHG (∼42 h post erythrocyte invasion) parasitized erythrocytes were exposed to tipranavir, lopinavir, ritonavir, saquinavir or DMSO control for 12 h before the preparation of Giemsa stained thin smears. Representative images of treatments are shown.

We next attempted to directly assess whether tipranavir, lopinavir, ritonavir or saquinavir can inhibit the processing of the plasmepsin IX [12] and plasmepsin X [12, 13] substrate *Pf*RAP1 and the *Pf*SUB1 substrate *Pf*MSP1. We treated schizont stage parasites (∼42 h post invasion) with inhibitory concentrations of each drug for 6 h and then harvested cells for Western blot. At concentrations where loss of merozoite invasion was evident with extended schizont treatment (**Figure 2E-H**) there was no evidence of full-length *Pf*MSP1 (**Figure 4A**) or *Pf*RAP1 (**Figure 4B**) accumulation as would be expected if the processing of these proteins was inhibited by the antiretroviral protease inhibitors. However the treatment of parasites with concentrations of antiretroviral protease inhibitors that imhibited merozoite invasion did appear to reduce the overall expression of *Pf*MSP1 and *Pf*RAP1 when compared to the *Pf*EXP2 loading control (**Figure 4C**). This suggested that extended protease inhibitor treatment may impact the overall health and normal development of merozoites.

**Figure 5:**
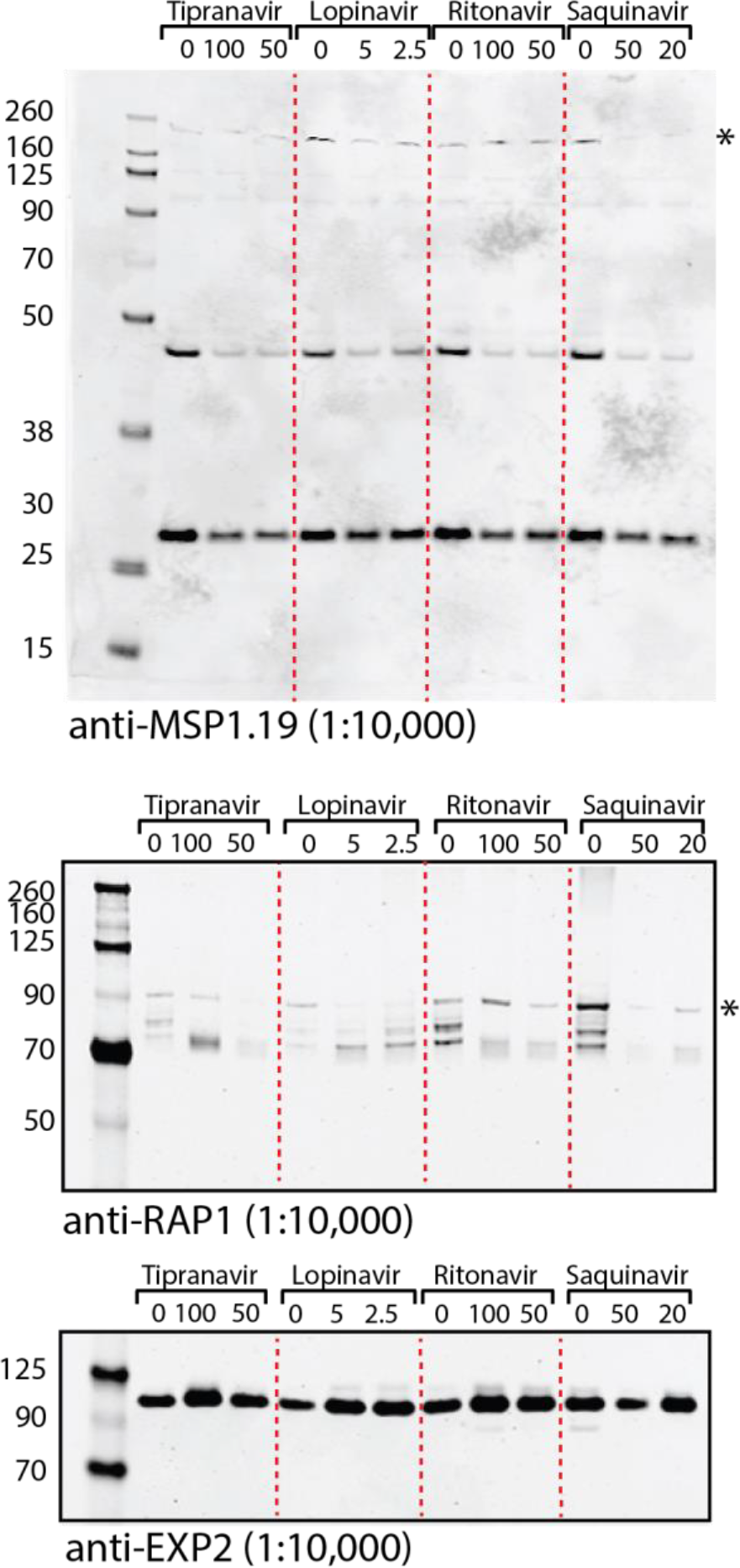
Effect of tipranavir, lopinavir, ritonavir, and saquinavir on *Pf*MSP1 and *Pf*RAP1 processing. Example Western blots showing *Pf*MSP1 **(A)** and *Pf*RAP1 **(B)** in *P. falciparum* following treatment with tipranavir, lopinavir, ritonavir or saquinavir. **(C)** EXP2 was used as a loading control. * indicates the expected size of full length *Pf*MSP1 and *Pf*RAP1.

## Discussion

Although antiretroviral protease inhibitors are attractive starting points to develop antimalarials due to their history of safety for treatment of HIV and broad efficacy against malaria parasites [1, 4-6], the antiplasmodial activity of these drugss is not understood. While multiple targets have been proposed by various groups [3, 7, 8], clear identification of a mode of action or specific target has remained elusive. Given their stage of expression and importance in parasite survival, plasmepsin IX and X were of interest as potential targets of the antiretroviral protease inhibitors. Recent progress in establishing that plasmepsin IX and X play essential roles in parasite erythrocyte egress and invasion [12, 13] suggested that testing the impact of antiretroviral protease inhibitors on these processes would provide more information on the action of these drugs against malaria parasites.

Our data describing the effects of tipranavir, lopinavir, ritonavir and saquinavir and on *P. falciparum* merozoite egress and invasion suggest that plasmepsin IX and X are unlikely to be the primary targets of these drugs. While all of the antiretroviral protease inhibitors tested inhibited the development and, or egress of merozoites from erythrocytes and subsequent merozoite invasion, they did so at concentrations above those determined to be effective agasint asexual parasites in *in vitro* assays [1, 3] (**Figure 1** and **2**). In addition, data from previous stage-specific activity experiments where synchonized ring, trophozoite and schizont parasites were exposed to 40 μM saquinavir or 20 μM lopinavir for 8 h (the same exposure time-frame used in egress and invasion experiments in this study), show that inhibition of parasite growth after trophozoite treatment (∼80% inhibition for 20 μM lopinavir) is more dramatic than the effect on egress and invasion (≤20% inhibition for 20 μM lopinavir; **Figure 2**). Similarly, the growth inhibitory activity of high concentration tipranavir against trophozoites (∼90% inhibition; 150 μM for 8 h) [5] was greater than the invasion and egress inhibitory activity of this drug (60-80%; 150 μM for 8 h; **Figure 2**). While the asexual stage IC_50_ values of tipranavir, ritonavir and saquinavir were higher in the current study than previously reported (**Figure 1**), the IC_50_ value of lopinavir was in line with previous reports [1, 3]. In addition Western blots of *Pf*RAP1 and *Pf*MSP1 processing in antiretrovial treated parasites suggested no inhibition of processing.

Taken together with previous studies that show selected antiretroviral protease inhibitors have an impact on multiple (non-egress or invasion related) biological targets within malaria parasites [2, 7], our data suggest that tipranavir, lopinavir, ritonavir and saquinavir are unlikely to have significant activity against plasmepsin IX and X. Given the differential activities of selected antiretroviral protease inhibitors against different biological processes and targets in malaria parasites, including glucose uptake [7] and haemoglobin digestion [3], we propose that each drug has multiple and different targets within malaria parasites. This idea fits well with published [19] and unpublished data generated in our laboratories which suggests that antiretroviral protease inhibitor resistance selection is difficult, requiring ≥ 9 months for 4-6 fold sensitivity changes, and that higher levels of resistance are associated with multiple genomic changes. While it is likely to be difficult to define the biological targets of the antiretroviral protease inhibitors against malaria parasites, the broad activity and safety of these drugs bodes well for their use and the use of similar compounds in the field.

## Declarations

### Ethics approval and consent to participate

Not applicable

### Consent for publication

Not applicable

### Availability of data and materials

The datasets used and/or analysed during the current study are available from the corresponding author on reasonable request

### Competing intersts

The authors declare that they have no competing interests to declare

### Author’s contributions

DWW and SF designed and performed all experiments. DWW and TSS-A analysed the data. All authors (DWW, SF, TSS-A and KTA) provided direction to the study, contributed to manuscript preparation and approved the final manuscript.

## Acknowledgements

We thank the Australian Red Cross Lifeblood^®^ for the provision of human blood for culture of *Plasmodium* parasites and AIDS Reagent Program, Division of AIDS, NIAID, NIH, for supplying tipranavir.

